# Flexible parental care: Uniparental incubation in biparentally incubating shorebirds

**DOI:** 10.1101/117028

**Authors:** Martin Bulla, Hanna Prüter, Hana Vitnerová, Wim Tijsen, Martin Sládeček, José A. Alves, Olivier Gilg, Bart Kempenaers

**Author notes:** Author for correspondence: Martin Bulla.

## Abstract

Recent findings suggest that relative investment of females and males into parental care depends on the population’s adult sex-ratio. For example, all else being equal, males should be the more caring sex if the sex ratio is male biased. Whether such outcomes are evolutionary fixed (i.e. related to the species’ typical sex-ratio) or whether they arise through flexible responses of individuals to the current population sex-ratio remains unclear. Nevertheless, a flexible response might be limited by evolutionary history when one sex loses the ability to care or when a single parent cannot successfully care. Here, we demonstrate that after the disappearance of one parent, individuals from 8 out of 15 biparentally incubating shorebird species were able to incubate uniparentally for 1-19 days (median = 3, *N* = 69). Such uniparental phases often resembled the incubation rhythm of species with obligatory uniparental incubation. Although it has been suggested that females of some shorebirds desert their brood after hatching, our findings indicate that either sex may desert prior to hatching. Strikingly, in 27% of uniparentally incubated clutches - from 5 species - we document successful hatching. Our data thus reveal the potential for a flexible switch from biparental to uniparental care.

## INTRODUCTION

Parental care is a tremendously diverse social trait. The extent of parental cooperation varies along a continuum, from equal share of care by both parents, to uniparental care in which either the female or the male provides all care^1,2^. Recent theoretical work and comparative empirical studies suggest that the sex that is in short supply in the population has increased mating opportunities, and is thus less likely to provide care than the more abundant sex^3–8^. Although these empirical studies provide some support for the role of the adult population sex-ratio in shaping parental care patterns on an evolutionary time-scale, it is less clear whether individuals can flexibly adjust their form of parental care in relation to the environment, including the current population sex-ratio. Essentially, the species’ evolutionary history might have fixed the pattern of parental care, leaving little room for flexibility in who cares.

In some species, the caring sex varies between pairs (e.g. ref. 9, 10–15). For example in some cichlid fish, males are more likely to desert their brood when opportunities to breed are high^9,10^. In several bird species, biparental care is facultative (e.g. ref. 13, 14–18) whereas in others it is considered obligatory^19,20^. Here, we focus on a specific form of avian parental care, incubation of eggs. In some species parents can switch flexibly between years from biparental to uniparental care and vice versa^14^. In others such flexibility is unlikely because, for example, one sex (often the male) lacks a brood patch and hence cannot incubate effectively^21^. Flexibility may also be limited in species where females and males possess a brood patch and share incubation roughly equally, because a single parent may not be able to attend the nest enough for embryos to develop until hatching, either because embryos cannot withstand fluctuating temperatures^19,22^, or because clutches that are left alone have a high probability to get predated^23^. On the other hand, flexibility might be favoured by selection, because it would allow a single individual to obtain at least some reproductive success when its partner disappears (e.g. because of predation or illness).

Here, we investigate the occurrence of uniparental incubation in 15 shorebird species (Table 1) that are considered ‘obligate’ biparental incubators^24–26^. First, we report the frequency of uniparental incubation and describe how daily nest attentiveness (incubation constancy) changed from a biparental to a uniparental rhythm. Second, we compare the uniparental incubation rhythms from biparental incubators with the incubation rhythms of obligatory uniparental shorebird species with female-only incubation (pectoral sandpiper, *Calidris melanotos*), as well as with male-only incubation (red-necked phalarope, *Phalaropus lobatus*). Finally, we investigate whether the uniparentally incubated clutches succeeded (i.e. at least one chick hatched) and, if so, whether hatching success was related to when the uniparental phase started within the incubation period, to the duration of the uniparental phase, and to the median daily nest attendance.

**Table 1.**
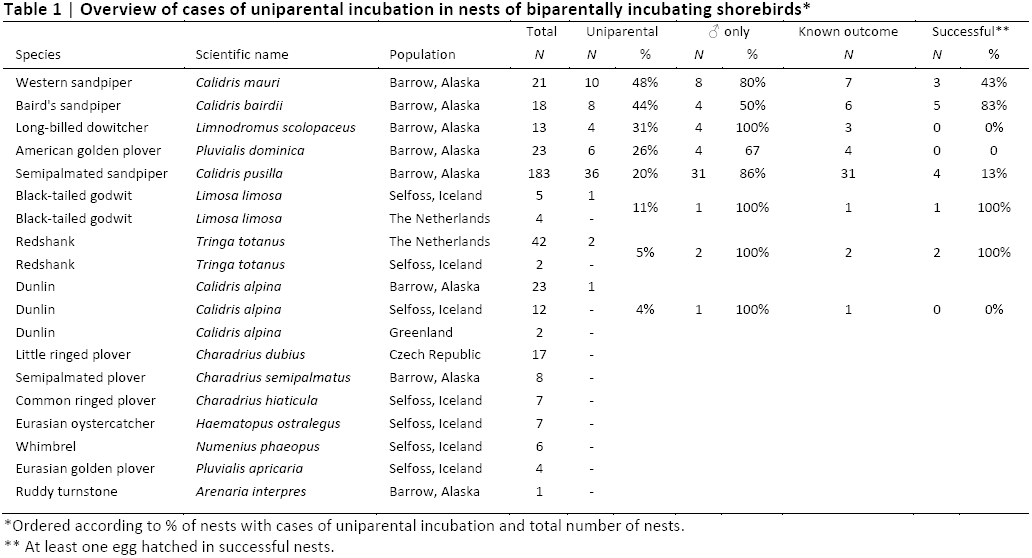
Overview of cases of uniparental incubation in nests of biparentally incubating shorebirds*.

## METHODS

### Data collection

Between 2011 and 2015, we recorded incubation with radio frequency identification (RFID) in combination with nest and surface (next to the nest) temperature probes^27,28^ at hundreds of nests from 19 populations of 15 biparentally incubating shorebird species (Table 1); eight populations were breeding in Alaska, seven in Iceland, one in the Czech Republic, one in Greenland, and two in the Netherlands (for information on the study sites see ref. 29, 30). In 2008 and 2009, we recorded uniparental incubation of 13 female pectoral sandpipers in Barrow, Alaska (71.32°N, 156.65°W) using an automated tracking system^31^. In 2015, we recorded uniparental incubation of 9 male red-necked phalaropes from Chukotka (64.75°N, 177.67°E) using nest and surface temperature probes^27,28^.

If birds were monitored already prior to laying, we estimated the start of incubation from the visualized raw data (actograms). If a nest was found during laying, we estimated the start of incubation by assuming that females laid one egg per day and started incubation when the clutch was completed (usually four, rarely three eggs). If nests were found with a full clutch, we estimated the start of incubation by subtracting the average incubation period of the species (derived from the literature, see Metadata in ref. 32) from the hatching date, or as the median start of incubation derived from the height and inclination of the eggs floated in water^33^. For one nest where we lacked the relevant information, we estimated the start of incubation as the median start of incubation of the species in that population and year.

Some of the studied nests were protected against avian predators using one of two enclosure types, both made of mesh wire (see Supplementary 1, Picture S1 in ref. 27 and Supplementary Fig. 1 in ref. 22).

All field procedures were performed in accordance with the relevant guidelines and regulations, and approved by the Max Planck Institute for Ornithology, and the US, Icelandic, Dutch, Czech and Russian authorities.

### Extraction of incubation behaviour

For each nest, we transformed the incubation records to local time as UTC time + / (longitude of the nest/15). In nests with temperature recordings, constant nest-temperatures above the surrounding surface temperatures were interpreted as continuous incubation; the start of incubation was determined from the steep increase in nest temperature, the end of incubation from a steep decrease in temperature^27, 28, 32, 34^.

For nests with automated tracking, we used changes in signal strength from a radio-tag attached to the rump of the female: incubation was inferred whenever signal strength remained nearly constant (for details see ref. 31 and Scripts in ref. 32).

### Definition of uniparental incubation

We report on 70 cases of uniparental incubation that occurred after a parent naturally disappeared (either deserted the nest or died, which was usually unknown; *N* = 54 cases), after a parent deserted following capture and release (*N* = 13 cases), or after we experimentally removed a parent (*N* = 3 cases from semipalmated sandpiper^22^).

When a parent disappears, its partner first incubates its regular incubation bout^22^, here defined as the median incubation bout length of that population (see Data in ref. 32, derived from ref. 30, 35). Then the ‘deserted’ parent often attempts to compensate for the ‘supposed bout’ of its partner^22^ and then it may immediately desert. We do not consider such transient cases as uniparental incubation.

We thus here define uniparental incubation as cases where a single parent incubated for at least twice the median incubation bout of the population, excluding the first regular incubation bout and the periods 6 hours before the start of hatching or 24 hours before the chicks had hatched or left the nest. In this way we limited the data to true uniparental incubation periods that did not include one prolonged incubation bout (due to the partner’s absence), cases where this bout was followed by immediate and complete nest desertion, and periods that were not confounded by hatching. Furthermore, including only longer periods of uniparental incubation allowed us to investigate the change in nest attendance within a day or over several days, i.e. from an initial ignorance about the partner’s absence or from an initial attempt to compensate for a possible delayed return of the partner^22^ to an individual’s response to the partner’s ‘desertion’.

In total, we identified 70 periods of uniparental incubation from 68 nests; two nests had two periods, because one of the parents first ‘deserted’ the nest temporarily, came back to incubate, and then ‘deserted’ again with no subsequent return.

### Definition of nest attendance

To compare incubation patterns between biparental and uniparental periods, as well as to compare uniparental incubation patterns between biparental and uniparental species, we use nest attendance (incubation constancy per hour or per day), defined as the proportion of time a bird actually incubated. For these analyses, we only included periods (a particular hour or a particular day) where at least 75% of the total time was either biparental or uniparental incubation. We also excluded one complete nest and part of the data from two nests, because the temperature readings failed due to a dislocated probe. We further excluded two nests where the uniparental bird incubated only a single egg. Thus, our data set on uniparental incubation included 895 data points for daily nest attendance and 23,258 data points for hourly nest attendance from a total of 87 nests from 10 species (65 nests of 8 biparental species, 22 nests of 2 uniparental species).

### Statistical analyses

#### Nest attendance

We tested for the difference between biparental and uniparental nest attendance in two mixed-effect models. The first model contained daily nest attendance (proportion) as response variable, and day in the incubation period - defined as the proportion of the species’ typical incubation period (available in Data of ref. 32, derived from ref. 24, 25) that had already passed – as a continuous predictor. We fitted incubation period in interaction with type of care (biparental or uniparental incubation in biparental species, uniparental incubation in uniparental species). Nest and species, both in interaction with type of incubation (biparental or uniparental), were included as random intercepts and day in the incubation period as random slope.

The second model contained hourly nest attendance (proportion) as response variable, and time of day (continuous; transformed to radians and represented by a sine and cosine) in interaction with type of care (three levels as above) as predictors. Nest and species, both in interaction with type of incubation (biparental or uniparental), were included as random intercepts, and time of day as random slope.

For four biparental species, where we observed both female and male uniparental incubation, we used two additional models to test whether uniparental incubation patterns differed between the sexes. The first model contained daily uniparental nest attendance as response variable, day in the incubation period (proportion of the species’ typical incubation period) in interaction with sex as predictors, nest and species as random intercepts and day in the incubation period as random slope. The second model contained hourly uniparental nest attendance as response variable, time of day (transformed to radians and represented by a sine and cosine) in interaction with sex as predictors, nest and species as random intercepts, and time of day as random slope.

#### Nest success

For 55 biparental nests with phases of uniparental incubation we had information about the fate of the nest and for 51 of those also information about nest attendance. Using a binomial model, we tested whether nest success (at least one egg hatched: yes or no) was related to when the uniparental incubation started within the incubation period (defined as above; referred to as ‘start’) and for how many days it lasted (duration), and to median daily nest attendance. Species was specified as random slope. The correlations between the three predictors were low (all |*r*_Pearson or Spearmen_|< 0.32, *N* = 50 nests).

#### General procedure

R version 3.3.0^36^ was used for all statistical analyses and the ‘lme4’ R package^37^ for fitting the mixed-effect models. We used the ‘sim’ function from the ‘arm’ R package and a non-informative prior-distribution^38,39^ to create a sample of 5,000 simulated values for each model parameter (i.e. posterior distribution). We report effect sizes and model predictions by the medians, and the uncertainty of the estimates and predictions by the Bayesian 95% credible intervals represented by 2.5 and 97.5 percentiles (95% CI) from the posterior distribution of 5,000 simulated or predicted values. We estimated the variance components using the ‘lmer’ or ‘glmer’ function from the ‘lme4’ R package^37^ with maximum likelihood.

## RESULTS

### Occurrence of uniparental incubation

We found at least one case of uniparental incubation in 8 out of the 15 biparental shorebird species that we studied (Table 1). Across species, the proportion of nests with uniparental incubation ranged from 4% to 48% (Table 1; median weighted by the total number of nests for a given species = 19.3%). Females incubated uniparentally less often than males (in 14 out of 70 cases, and in 4 out of 8 species; Figure 1a and Supplementary Table 1).

**Figure 1.**
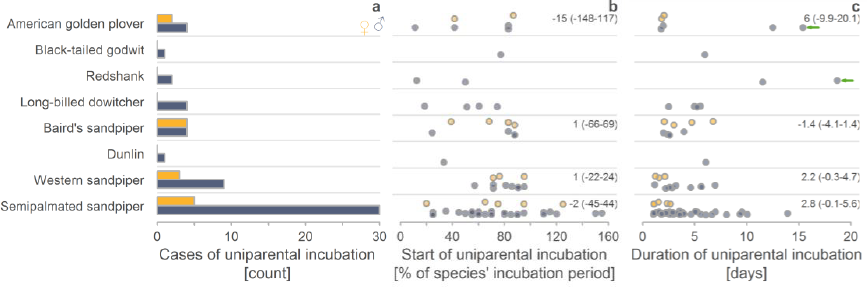
Uniparental incubation in eight biparental shorebirds according to sex. a, Frequency of uniparental incubation by females and males (*N* = 70 cases from 68 nests). The species are ordered by phylogeny. **b**, Distribution of the ‘start of uniparental incubation’ within the incubation period, defined as the % of the species’ typical incubation period that has already passed (*N* = 69 cases from 68 nests). Values larger than 100% indicate uniparental incubation that started after the typical species-specific incubation period had passed without hatching (see Methods). **c**, Distribution of the ‘duration of uniparental incubation’ in days (*N* = 69 cases from 68 nests). Green arrows indicate two nests where the male likely incubated uniparentally for the entire incubation period. **a-c**, Female uniparental incubation (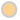 yellow), male uniparental incubation (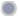; blue-grey). Data points are jittered to increase visibility. **b,c**, For species with cases of both male and female uniparental incubation, we give the posterior estimates (medians) of the effect sizes and the 95% credible intervals (CI) from a posterior distribution of 5,000 simulated values generated by the ‘sim’ function in R^39^ (based on a separate linear model for each species with sex as predictor variable).

Uniparental incubation started mostly in the second half of the incubation period (median = 71% of incubation period, range: 11 - 155%, *N* = 69 cases with known start of uniparental incubation from 68 nests of 8 species; Figure 1b). The median remained similar (70%) after we excluded uniparental incubation that started after the eggs were supposed to hatch (range: 11 – 95%, *N* = 62 cases from 60 nests of 8 species). Overall, the start of uniparental incubation within the incubation period was independent of sex (males differed from females by - 5.7%, CI: -31% - 20%, *N* = 69). Species (random slope) explained only 7% of the phenotypic variance (for estimates of sex differences for each species see Fig. 1b).

Uniparental incubation lasted a median of 3 days (range: 1 – 19 days, *N* = 69 cases; Figure 1c), which is an underestimation, because in 10 nests we removed the monitoring system before incubation ended and in three nests a single parent incubated ever since we found the nests. In two nests (one American golden plover *Pluvialis dominica* and one Common redshank *Tringa totanus*), a single parent probably incubated for the entire incubation period (see green arrow in Fig. 1c, and Supplementary Actograms^32^). Overall, uniparental incubation by males lasted 2.4 days longer than by females (CI: 0.5-4.3 days; *N* = 69 cases). However, species varied greatly in this respect (random slope explained 47% of the phenotypic variance; for estimates of sex differences for each species see Fig. 1c).

### Nest attendance during biparental and uniparental incubation

After the switch from biparental to uniparental incubation, daily nest attendance decreased and was overall similar to the daily nest attendance observed in uniparental species (Figure 2a, b & 3). Whereas daily nest attendance was constant over the incubation period in uniparental species and during biparental incubation (Figure 2b), daily nest attendance tended to increase over the incubation period during uniparental incubation in biparental species (Figure 2b and Supplementary Table 2). However, individuals varied greatly in this respect (random slope explained 35% of the variance, Supplementary Table 2), and for uniparentally incubating females of biparental species nest attendance seemed to decrease with incubation period (Supplementary Fig. 1 and Supplementary Table 3).

**Figure 2.**
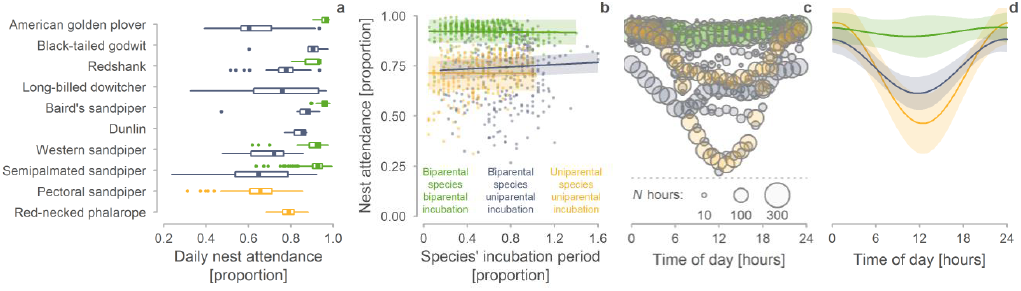
Daily nest attendance in biparental and uniparental shorebirds. a, Distribution of biparental and uniparental daily nest attendance. Box plots depict median (vertical thick line inside the box), the 25^th^ and 75^th^ percentiles (box), the 25^th^ and 75^th^ percentiles ±1.5 times the interquartile range or the minimum/maximum value, whichever is smaller (bars), and the outliers (dots). **b**, Change in daily nest attendance across the incubation period expressed as the proportion of the species’ typical incubation period. Each dot represents the nest attendance during one day. **c, d**, Change in hourly nest attendance across the day. Dots (**c**) represent mean hourly observations for each species (dot size reflects sample size). **b, d**, Lines with shaded areas indicate model predictions with 95%CI (Supplementary Table 2 & 4) based on the joint posterior distribution of 5,000 simulated values generated by the ‘sim’ function in R ^39^. **a-d,** Only nests that contain a uniparental incubation phase are included. Green 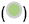 indicates biparental species during biparental phase, blue-grey 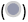 biparental species during uniparental phase, and yellow 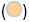 uniparental species. *N*_a-b_ = 895 days and *N*_c-d_ = 23,258 hours from 87 nests of 10 species (65 nests of 8 biparental species, 22 nests of 2 uniparental species).

Individuals of biparental species varied strongly in how they incubated uniparentally over the day. Some individuals continued to incubate as if their partner was still present, that is, they only incubated during ‘their’ bouts and left the nest unattended for the rest of the day when their partner was ‘supposed to’ incubate (e.g. actograms biparental_33, 38, 42, & 51 in Supplementary Actograms^32^). Other individuals developed an incubation rhythm similar to that of uniparental species, with continuous nest attendance at ‘night’ when it is colder and intermittent incubation – with short feeding bouts – during the day when it is warmer (e.g. biparental_15, 70, 73, 76-7^32^). However, within-individual variation in hourly nest attendance was far greater than the between-individual variation (within-individual [residual variance] = 53% and between-individual = 8% of variance; Supplementary Table 4). Indeed, some individuals first incubated as if their partner was still present and then switched to a ‘uniparental-like’ rhythm (e.g. biparental_26, 35, 42^32^). When individuals continued to incubate as if their partner was still present nest attendance was about 10-20% lower than when individuals incubated like uniparental species (Supplementary Actograms^32^). One male redshank kept a ‘uniparental like’ rhythm for 18 days (biparental_77^32^), and one male semipalmated sandpiper for about 10 days (biparental_37^32^).

The 24-hour rhythm of uniparental incubation in biparental species often closely resembled that of uniparental species with high nest attendance at night when it is colder and low nest attendance during the day when it is warmer (Figure 3 and Figure 2c, d; for nest-specific relationships see Supplementary Fig. 2). In contrast, during biparental incubation, nest attendance was always high, with only a slight dip during the warmest part of the day (Figure 2c, d). The rhythm of uniparental nest attendance in biparental species was similar for females and males (Supplementary Fig. 1, Supplementary Table 5).

**Figure 3.**
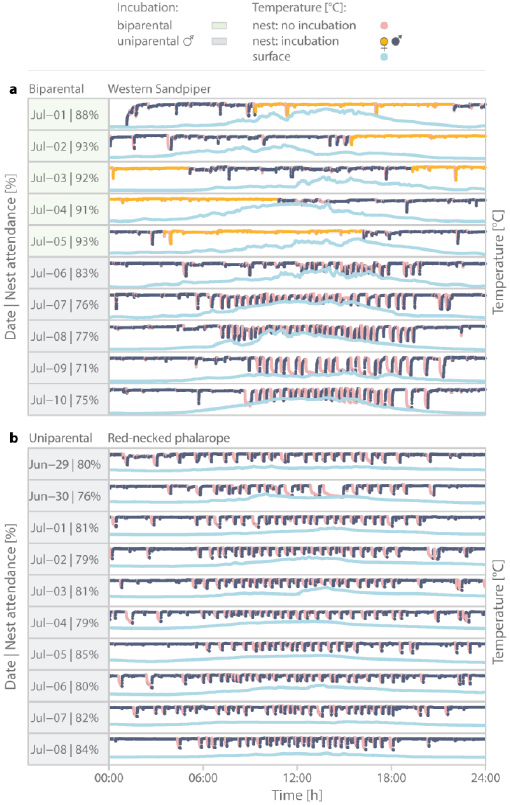
Example of a uniparental incubation rhythm by a biparental and a uniparental shorebird. a, Biparental shorebird (Western sandpiper) with a switch from biparental incubation (days marked green, 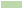) to uniparental male-only incubation (grey, 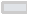). **b**, Uniparental species (Red-necked phalarope) with male-only incubation. **a, b**, Pink 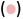 demarcates nest temperatures considered as no incubation. Yellow 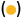 demarcates nest temperatures considered as incubation while female was on the nest and dark-blue 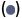 while male was on the nest. Light-blue 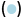 indicates surface temperatures near the nest. Temperatures were recorded every 5 s. Daily nest attendance is defined as the percentage of incubation readings (yellow and dark-blue; 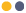) from all nest temperature readings for that day (pink + yellow + dark-blue; 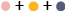).

### Nest success for biparental species under uniparental incubation

Out of 55 uniparentally incubated nests (from 8 species) for which we knew the outcome, at least one chick hatched in 15 nests (27%; representing 5 species; Table 1), 4 nests (7%) were depredated and the remaining 36 nests (65%) were deserted. The nest success was independent of how uniparental incubation started (Supplementary Table 6), was different among species (ranged from 0-100%; Table 1), and in 5 out of 8 species was lower than under biparental incubation (Figure 4). However, the sample sizes for uniparental incubation are typically small (Table 1).

**Figure 4.**
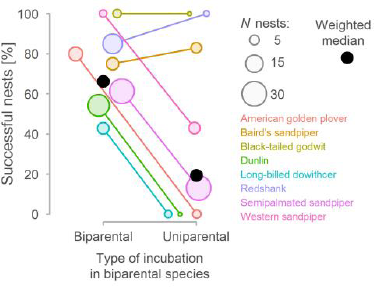
Nest success in nests of biparental species with biparental and uniparental incubation. Circles represent species. Colour indicates and lines connect same species. Black dots indicate median, weighted by sample size (reflected in the size of the circles). The data for biparental species come from the same populations as the uniparental data and were extracted from ref. 30, 35.

The successful nests became uniparental later in the incubation period and were incubated uniparentally more day and with higher median nest attendance than the nests that failed (Figure 5 and Supplementary Fig. 3). Parents could successfully hatch their eggs even if they continued uniparental incubation after the ‘normal’ incubation period had ended (Supplementary Fig. 3). However, singly-incubating parents were unsuccessful if they started uniparental incubation after the ‘normal’ incubation period had ended (Figure 5a and Supplementary Fig. 3), suggesting that lack of hatching at the expected date may lead one parent to desert.

**Figure 5.**
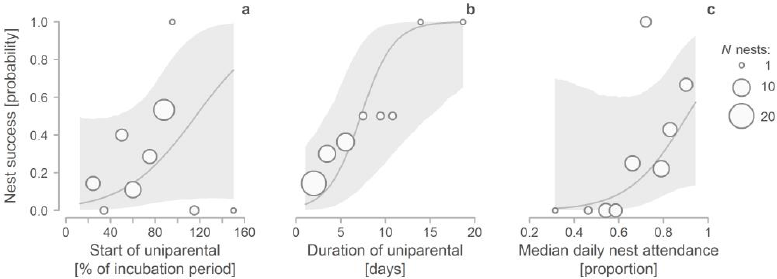
Predictors of nest success (hatching of at least one egg) for biparental species under uniparental incubation. a-c, Probability of nest success as a function of when the uniparental incubation started within the incubation period (expressed as the % of the species’ incubation period that had passed when uniparental incubation started; **a**), as a function of how long the uniparental incubation lasted (**b**), and as a function of median daily nest attendance (**c**). Circles represent means for intervals spread evenly across the range of x-values; circle size reflects sample size. The solid lines depict the model-predicted relationships, the shaded area the 95% credible intervals based on the joint posterior distribution of 5,000 simulated values generated by the ‘sim’ function in R^39^; the predicted relationships stem from a binomial mixed-effect model (Supplementary Table 7), where the effect of the other predictors was kept constant. *N* = 50 nests with uniparental incubation from 8 biparental species.

## DISCUSSION

Our findings reveal that phases of uniparental incubation are not uncommon in biparental shorebirds, and challenge the belief that this necessarily leads to nest failure^25^. We found uniparental incubation in 8 out of 15 biparentally incubating shorebird species, and evidence of successful hatching in 5 of these species, that is in 27% of all uniparentally incubated nests with known outcome (Table 1). Reports of successful single-parent incubation from other species with ‘obligatory’ biparental incubation are rare^40^. Successful uniparental incubation in biparental incubators might truly be rare, but its frequency might be underestimated, because records of incubating parents throughout the entire incubation period are scarce^28^.

In biparental shorebirds, females typically desert their brood after hatching^40^. Here, we describe 68 cases (17% of nests) where one parent disappeared prior to hatching, and indeed it was more often the female (80% of nests). In most of these nests, desertion is likely, but for two nests our video recordings revealed that one of the parents had been taken by a predator. We cannot exclude that this also occurred in other nests. Furthermore, when uniparental incubation occurs closer to hatching, we cannot exclude the possibility that the ‘deserting’ parent left to replenish its energy stores and later re-joined its partner to brood and guide the chicks.

We found substantial variation in when one of the two parents deserted or disappeared from the nest and for how long the nest was incubated uniparentally (Figure 1b, c & 4b, c). In 10% of cases uniparental incubation only started after the (typical) incubation period of the species had ended (such nests always failed). In other cases, individuals incubated uniparentally for at least half and up to nearly the entire incubation period (Figure 1c). However, uniparental incubation typically started when 70% of the incubation period had passed (Figure 1b & 4b) and lasted about three days (Figure 1c & 4c). This suggests that individuals either continued incubating for a few days before deserting the nest, perhaps once realizing that they were incubating alone (which is in line with our experimental findings^22^), or incubated uniparentally only for few days until the eggs started hatching.

Importantly, we found that uniparentally incubated clutches from biparental species can successfully hatch, in particular when uniparental incubation started later in the incubation period (but before expected hatching), when uniparental incubation lasted longer (i.e. the parent did not give up) and when nest attendance by the single parent was higher (Figure 5). This suggests that in these biparental species one of the parents might benefit from deserting the nest, at least under some conditions, leaving the remainder of parental care to the partner. Indeed, some single parents were able or willing to incubate with a rhythm that closely resembled that of uniparental species (Figure 3).

If individuals from what are considered obligatory biparental species can behave as a uniparental species - with continuous incubation during the colder night and intermittent incubation with short foraging bouts during the day - then the potential exists for a flexible switch from biparental to uniparental care. Such flexibility may then lead to facultative biparental care^14, 18, 41^ or even to reduced or no care in one of the sexes. In turn, this could lead to a more flexible mating system including social polyandry and social polygyny^16, 42^. Our results reveal that male uniparental incubation was more common than female uniparental incubation (Figure 1). Thus, all else being equal, the evolution of polyandry would be more likely than the evolution of polygyny. It is worth investigating (a) whether flexible switches from biparental to (full) uniparental care occur in response to changing conditions (e.g. in response to warmer climate or to changes in mate availability), and (b) which factors determine who cares (e.g. population sex ratio).

### Open data, codes and materials

All available at https://osf.io/3rsny^32^.

### Authors’ contributions

M.B. and B.K. conceived the study; M.B. with help of B.K., and H.P. collected the biparental data in Barrow. H.V. and M.B. with help of B.K. and J.A.A. collected the data in Iceland; J.A.A. collected the Iceland godwit data and part of the oystercatcher data, O.G. collected the data on dunlins from Greenland. H.V. and M.B. collected the data in the Czech Republic. W.T. and H.P. collected the redshank data in Holland, M.B. collected the godwit data in Friesland, B.K. collected the pectoral sandpiper data, M.S. the phalarope data; M.B. coordinated the study, analysed the data, prepared all supporting information available from Open Science Framework^32^ and drafted the manuscript. All authors commented on the draft and M.B. with help of B.K. finalized the paper.

## Acknowledgements

We thank A. Rutten, E. Buchel, E. Stich, E. Vozabulová, F. Heim, F. Prüter, I. Steenbergen, J. Mlíkovský, K. Chmel, K. Murböck, L. Langlois, L. Verlinden, M. Schneider, M. Šálek, M. Valcu, S. Herber, S. Hobma, V. Dvořák, V. Heuacker, and V. Kubelka for help in the field, A. Girg for the genetic sexing, M. Valcu for advice on data anlysis, D. Starr-Glass and B. Bulla for constructive suggestions on the manuscript. M.B. thanks Bare and Maje for patience and support. M.B. did this work as a PhD student in the International Max Planck Research School for Organismal Biology.

## Competing interest

We have no competing interests.

## Funding

This work was funded by the Max Planck Society (to B.K.), EU Marie Curie individual fellowship at NIOZ Royal Netherlands Institute for Sea Research, Department of Coastal Systems (to M.B.), Ministry of Education, Youth and Sports of the Czech Republic (Grant MŠMT Kontakt II, project LH13278 “Interspecific relationships and predation risks in grassland and wetland bird communities”; to M.S.), FCT (SFRH/BPD/91527/2012; to J.A.A.), RANNIS (130412-051; to J.A.A.) and French Polar Institute IPEV Program 1036 (to O.G.).

## SUPPLEMENTARY INFORMATION

**Supplementary Table 1.** Cases of uniparental incubation according to sex and desertion circumstances.

| Desertion | Uniparental incubation |  |
| --- | --- | --- |
|  | ♀ | ♂ |
| After catching | 2 | 11 |
| Unknown reason | 10 | 41 |
| Removal experiment | 0 | 3 |
| Temporal | 2 | 1 |

**Supplementary Table 2.**
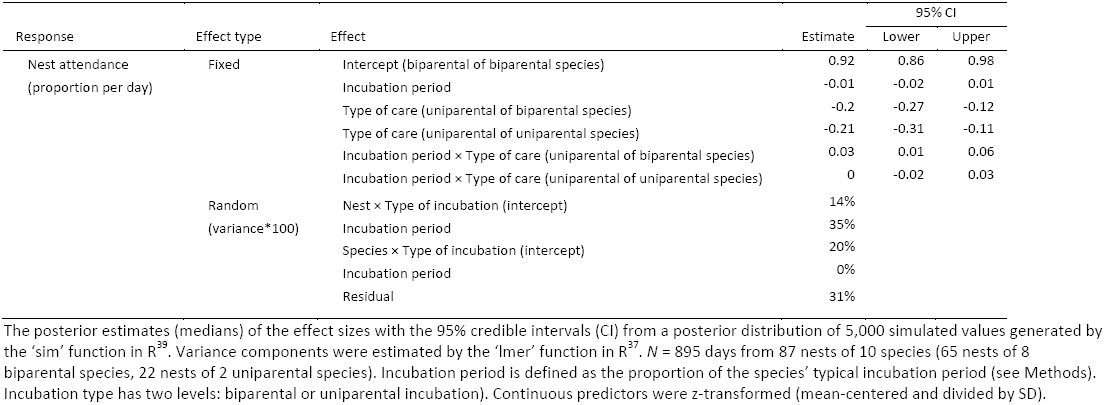
Effects of incubation type and incubation period on daily nest attendance.

| Response | Effect type | Effect | Estimate | 95% CI |  |
| --- | --- | --- | --- | --- | --- |
|  |  |  |  | Lower | Upper |
| Nest attendance<br>(proportion per day) | Fixed | Intercept (biparental of biparental species) | 0.92 | 0.86 | 0.98 |
|  |  | Incubation period | -0.01 | -0.02 | 0.01 |
|  |  | Type of care (uniparental of biparental species) | -0.2 | -0.27 | -0.12 |
|  |  | Type of care (uniparental of uniparental species) | -0.21 | -0.31 | -0.11 |
|  |  | Incubation period × Type of care (uniparental of biparental species) | 0.03 | 0.01 | 0.06 |
|  |  | Incubation period × Type of care (uniparental of uniparental species) | 0 | -0.02 | 0.03 |
|  | Random<br>(variance*100) | Nest × Type of incubation (intercept) | 14% |  |  |
|  |  | Incubation period | 35% |  |  |
|  |  | Species × Type of incubation (intercept) | 20% |  |  |
|  |  | Incubation period | 0% |  |  |
|  |  | Residual | 31% |  |  |

The posterior estimates (medians) of the effect sizes with the 95% credible intervals (CI) from a posterior distribution of 5,000 simulated values generated by the 'sim' function in R<sup>39</sup>. Variance components were estimated by the 'lmer' function in R<sup>37</sup>. $N = 895$ days from 87 nests of 10 species (65 nests of 8 biparental species, 22 nests of 2 uniparental species). Incubation period is defined as the proportion of the species' typical incubation period (see Methods). Incubation type has two levels: biparental or uniparental incubation). Continuous predictors were z-transformed (mean-centered and divided by SD).

**Supplementary Table 3.** Effects of sex and incubation period on daily nest attendance for biparental species during uniparental incubation.

| Response | Effect type | Effect | Estimate | 95% CI |  |
| --- | --- | --- | --- | --- | --- |
|  |  |  |  | Lower | Upper |
| Nest attendance<br>(proportion per day) | Fixed | Intercept (♀) | 0.75 | 0.66 | 0.84 |
|  |  | Incubation period | -0.05 | -0.12 | 0.03 |
|  |  | Sex (♂) | -0.06 | -0.13 | 0.02 |
|  |  | Incubation period × Sex (♂) | 0.1 | 0.02 | 0.17 |
|  | Random<br>(variance*100) | Nest (intercept) | 34% |  |  |
|  |  | Incubation period | 5% |  |  |
|  |  | Species (intercept) | 2% |  |  |
|  |  | Incubation period | 10% |  |  |
|  |  | Residual | 59% |  |  |

The posterior estimates (medians) of the effect sizes with the 95% credible intervals (CI) from a posterior distribution of 5,000 simulated values generated by the 'sim' function in R<sup>39</sup>. Variance components were estimated by the 'lmer' function in R<sup>37</sup>. $N = 218$ days from 56 nests of 4 species. Incubation period is defined as the proportion of the species' typical incubation period (see Methods). Continuous predictors were z-transformed (mean-centered and divided by SD).

**Supplementary Table 4.** Effects of incubation type and time of day on hourly nest attendance.

| Response | Effect type | Effect | Estimate | 95% CI |  |
| --- | --- | --- | --- | --- | --- |
|  |  |  |  | Lower | Upper |
| Nest attendance<br>(proportion per hour) | Fixed | Intercept (biparental of biparental species) | 0.92 | 0.86 | 0.98 |
|  |  | sin (time) | -0.01 | -0 | 0.03 |
|  |  | cos (time) | 0.02 | -0.1 | 0.09 |
|  |  | Type of care (uniparental of biparental species) | -0.17 | -0.3 | -0.1 |
|  |  | Type of care (uniparental of uniparental species) | -0.2 | -0.3 | -0.1 |
|  |  | sin (time) × Type of care (uniparental of biparental species) | -0 | -0 | 0.04 |
|  |  | sin (time) × Type of care (uniparental of uniparental species) | 0.04 | -0 | 0.09 |
|  |  | cos (time) × Type of care (uniparental of biparental species) | 0.11 | 0.02 | 0.2 |
|  |  | cos (time) × Type of care (uniparental of uniparental species) | 0.23 | 0.11 | 0.35 |
|  | Random<br>(variance*100) | Nest × Type of incubation (intercept) | 8% |  |  |
|  |  | sin (time) | 15% |  |  |
|  |  | cos (time) | 14% |  |  |
|  |  | Species × Type of incubation (intercept) | 4% |  |  |
|  |  | sin (time) | 0% |  |  |
|  |  | cos (time) | 6% |  |  |
|  |  | Residual | 53% |  |  |

The posterior estimates (medians) of the effect sizes with the 95% credible intervals (CI) from a posterior distribution of 5,000 simulated values generated by the 'sim' function in R<sup>39</sup>. Variance components were estimated by the 'lmer' function in R<sup>37</sup>. $N = 23,258$ hours from 87 nests of 10 species (65 nests of 8 biparental species, 22 nests of 2 uniparental species). Incubation type has two levels (biparental or uniparental incubation).

**Supplementary Table 5.**
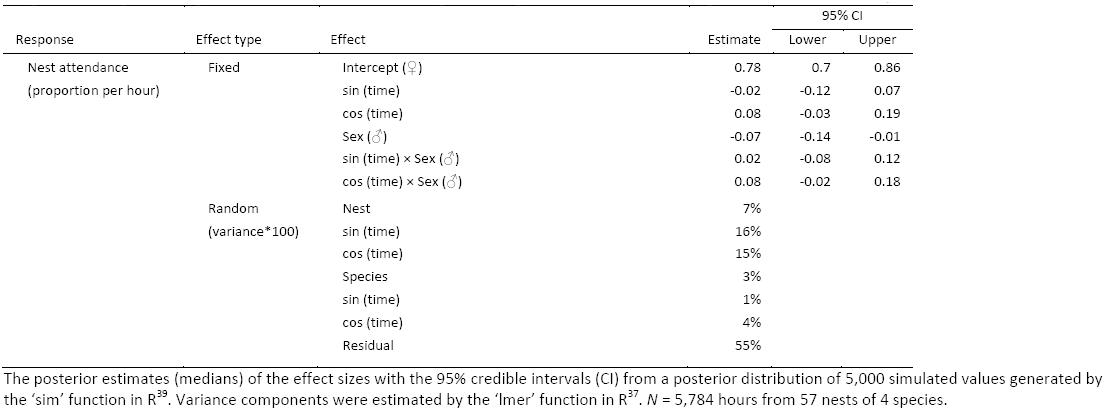
Effects of sex and time of day on hourly nest attendance for biparental species during uniparental incubation.

**Supplementary Table 6.** Cases of uniparental incubation according to desertion circumstances and nest faith*.

| Desertion | Nest successful |  |
| --- | --- | --- |
|  | Yes | No |
| After catching | 3 | 8 |
| Unknown reason | 12 | 29 |
| Removal experiment | 0 | 3 |
| Temporal | 2 | 1 |

\*In successful nests at least one egg hatched.

**Supplementary Table 7.** Effects of start and duration of uniparental incubation on the probability of hatching*.

| Response | Effect type | Effect | Estimate | 95% CI |  |
| --- | --- | --- | --- | --- | --- |
|  |  |  |  | Lower | Upper |
| Nest success<br>(1,0; on binomial scale) | Fixed | Intercept | -1.54 | -3.59 | 0.45 |
|  |  | Start (% of incubation period) | 0.93 | -0.32 | 2.26 |
|  |  | Duration (days) | 1.92 | 0.49 | 3.34 |
|  |  | Daily nest attendance (proportion) | 1.11 | -0.3 | 2.58 |
|  | Random<br>(variance) | Species (intercept) | 2.69 |  |  |

The posterior estimates (medians) of the effect sizes with the 95% credible intervals (CI) from a posterior distribution of 5,000 simulated values generated by the 'sim' function in R<sup>39</sup>. Variance components were estimated by the 'lmer' function in R<sup>37</sup>. $N = 50$ nests of 8 species. 'Start' defines when the uniparental incubation started within the incubation period (expressed as the % of the species' incubation period that had passed when uniparental incubation started). 'Duration' defines for how many days the uniparental incubation lasted and 'daily nest attendance' defined as median daily nest attendance. The predictors were z-transformed (mean-centered and divided by SD). The estimates are on a binomial scale: at least one egg hatched or not. Note that ideally a random slope should be added, but such model fails to converge because of insufficient data for each species.
\*In successful nests at least one egg hatched.

**Supplementary Figure 1.**
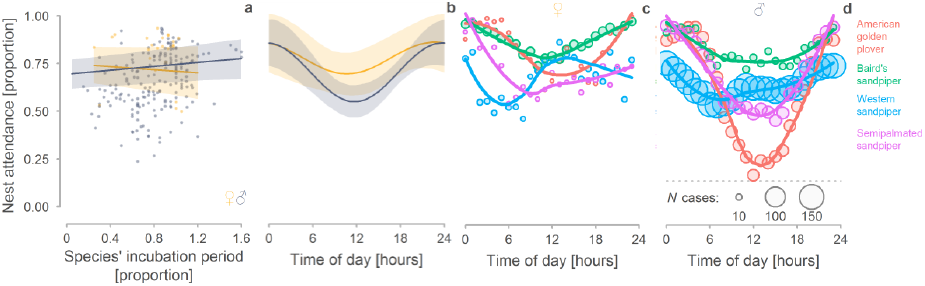
Sex-specific nest-attendance by four biparental species during uniparental incubation. a, Changes in daily nest attendance of uniparentally incubating females (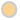; yellow) and males (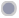; blue-grey) across the incubation period, expressed as the proportion of the species’ typical incubation period. Each dot represents the nest attendance during one day (*N* = 215 days for 56 nests of 4 species). **b-d,** Changes in hourly nest attendance across the day (*N*= 5,784 hours from 57 nests of 4 species). **b,** Lines with shaded areas indicate model predictions with 95% CI (Supplementary Table 3 & 5) based on the joint posterior distribution of 5,000 simulated values generated by the ‘sim’ function in R^39^. Colours as in (**a**). **c, d,** Dots represent mean hourly nest attendance for females (**c**) and males (**d**) per species (dot size reflects sample size,). Lines represent locally weighted scatterplot smoothing for each species.

**Supplementary Figure 2.**
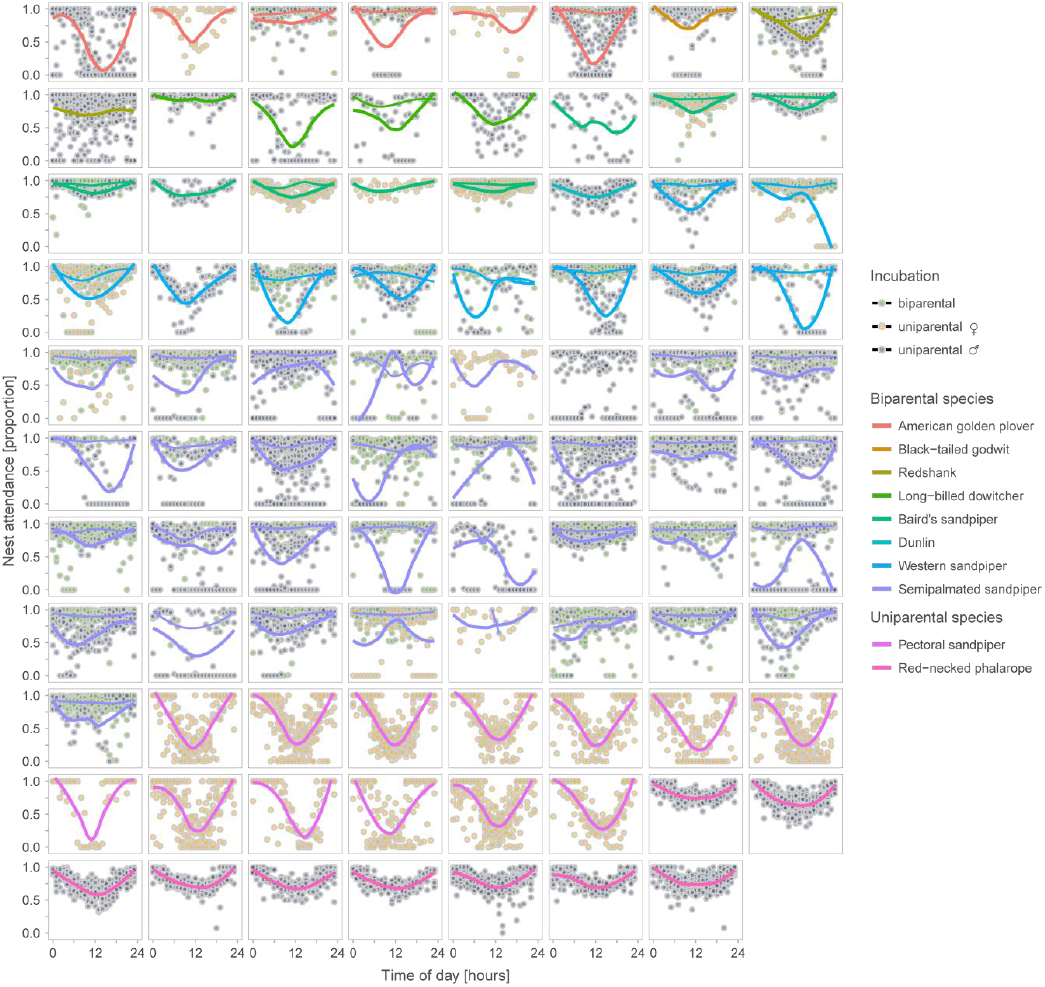
Variation in daily nest attendance. Each panel represents one of 87 nests. Dots represent hourly nest attendance, green dots 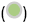 biparental incubation, yellow dots 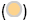 female uniparental incubation, and blue-grey dots 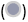 male uniparental incubation (note that green dots are often over plotted by the blue or yellow dots). Lines represent locally weighted scatterplot smoothing for each nest and incubation type (generated by the ‘ggplot2’ R package^43^). Line colour indicates species; thin lines 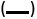 represent biparental incubation (only in nests with at least a day of monitored biparental incubation), thick lines 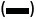 uniparental incubation.

**Supplementary Figure 3.**
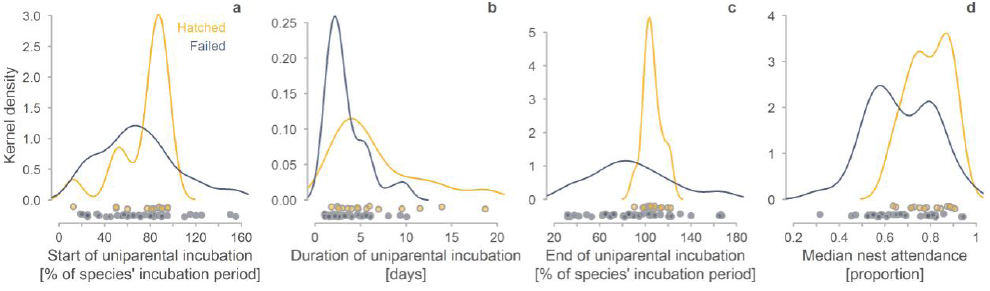
Distribution of descriptors of uniparental incubation from biparental species according to nest faith. a-d, Distribution of the start of uniparental incubation (expressed as the % of the species’ incubation period that had passed when uniparental incubation started, **a**), of the duration of uniparental incubation (**b**), of the end of uniparental incubation (expressed as the % of the species’ incubation period that had passed when uniparental incubation ended, **c**), and of the median daily nest attendance (**d**) in successful nests (at least one egg hatched; 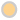, yellow; *N* = 15) and failed nests (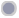, blue-grey; *N*_a,b_ = 39, *N*_c_ = 40, *N*_a,b_ = 36).

